## Supplementary figures and images for "Nanopore Guided Annotation of Transcriptome Architectures"

### Figure S1

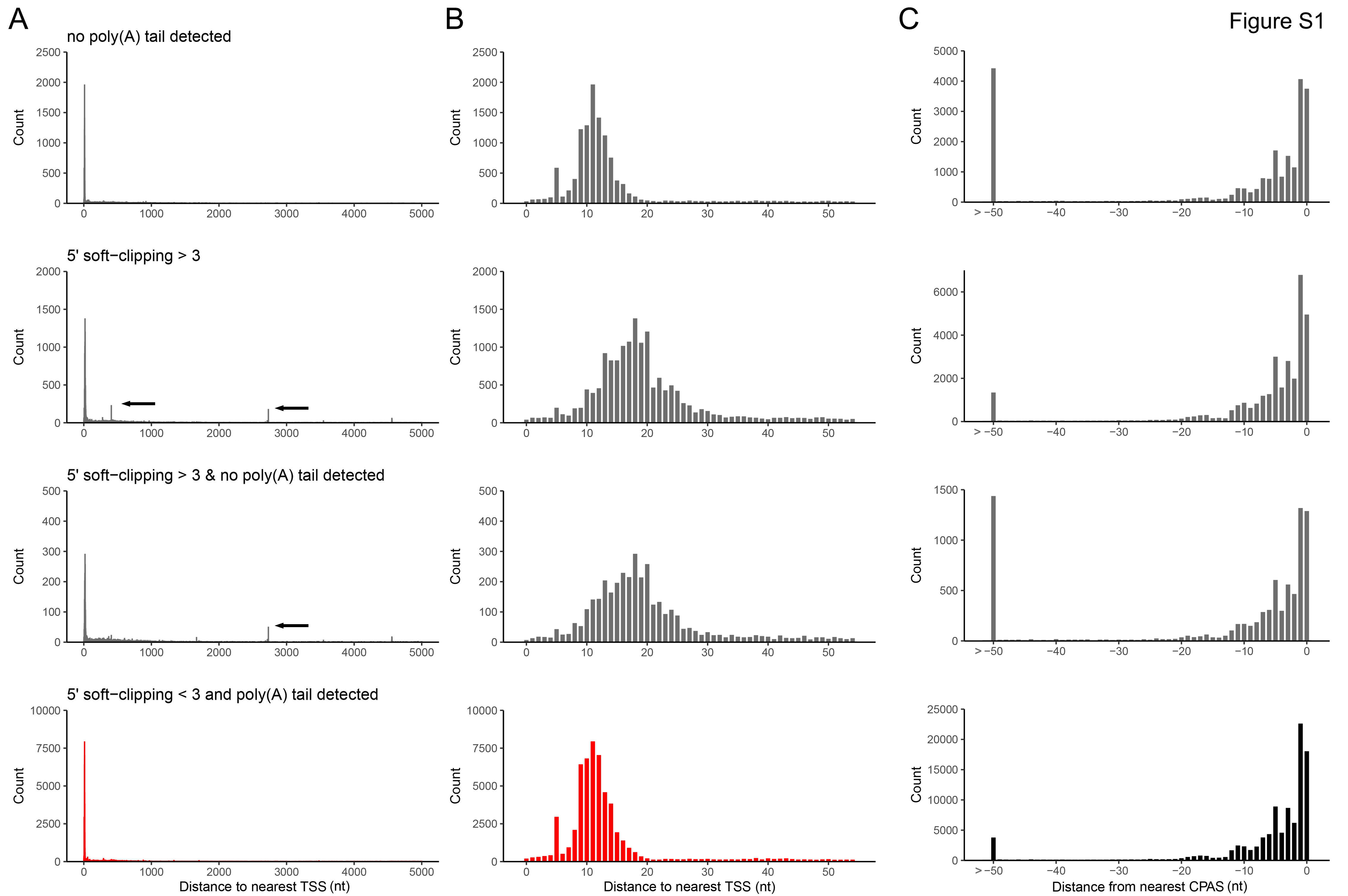

### Figure S2

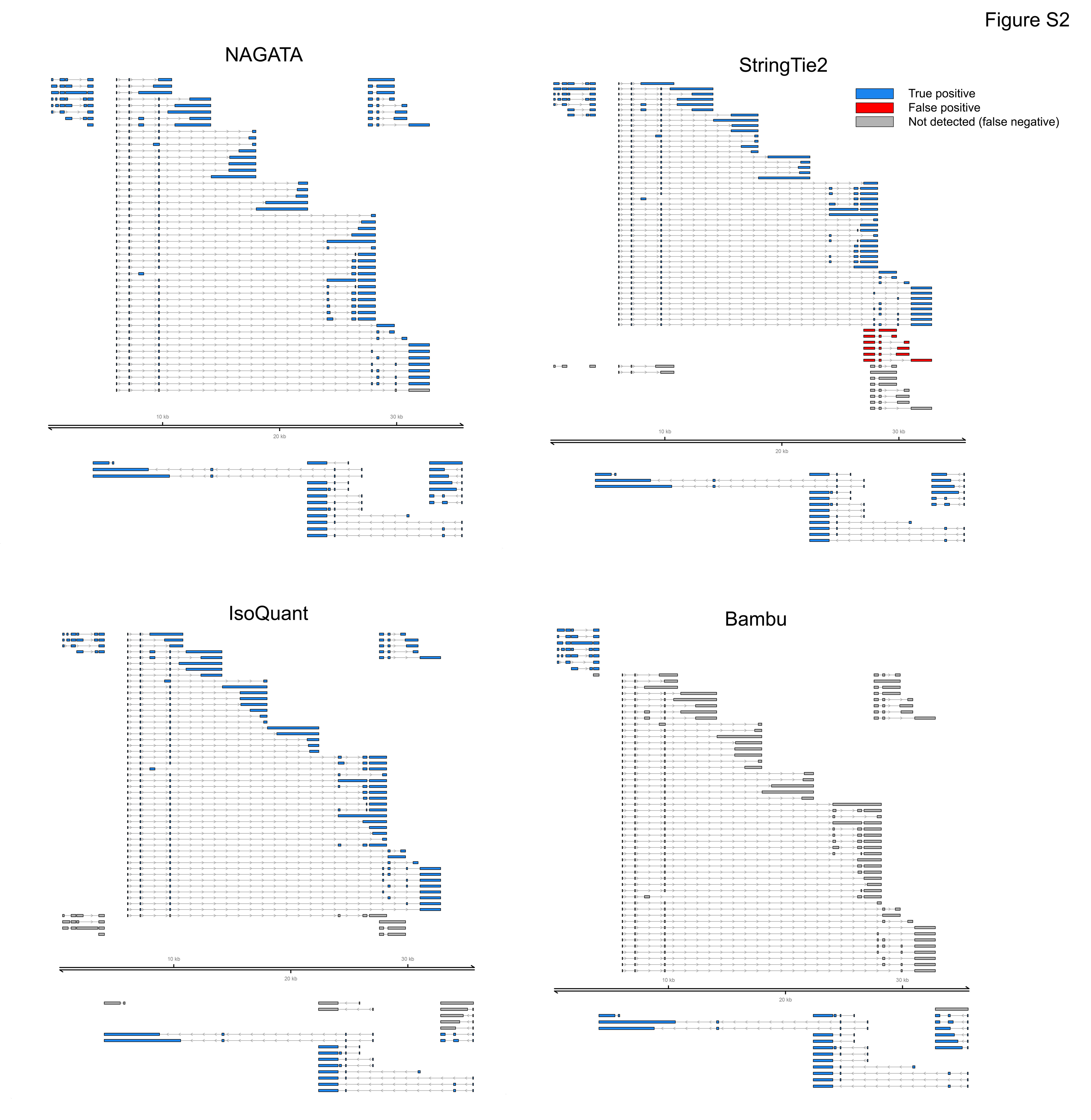

### Figure S3

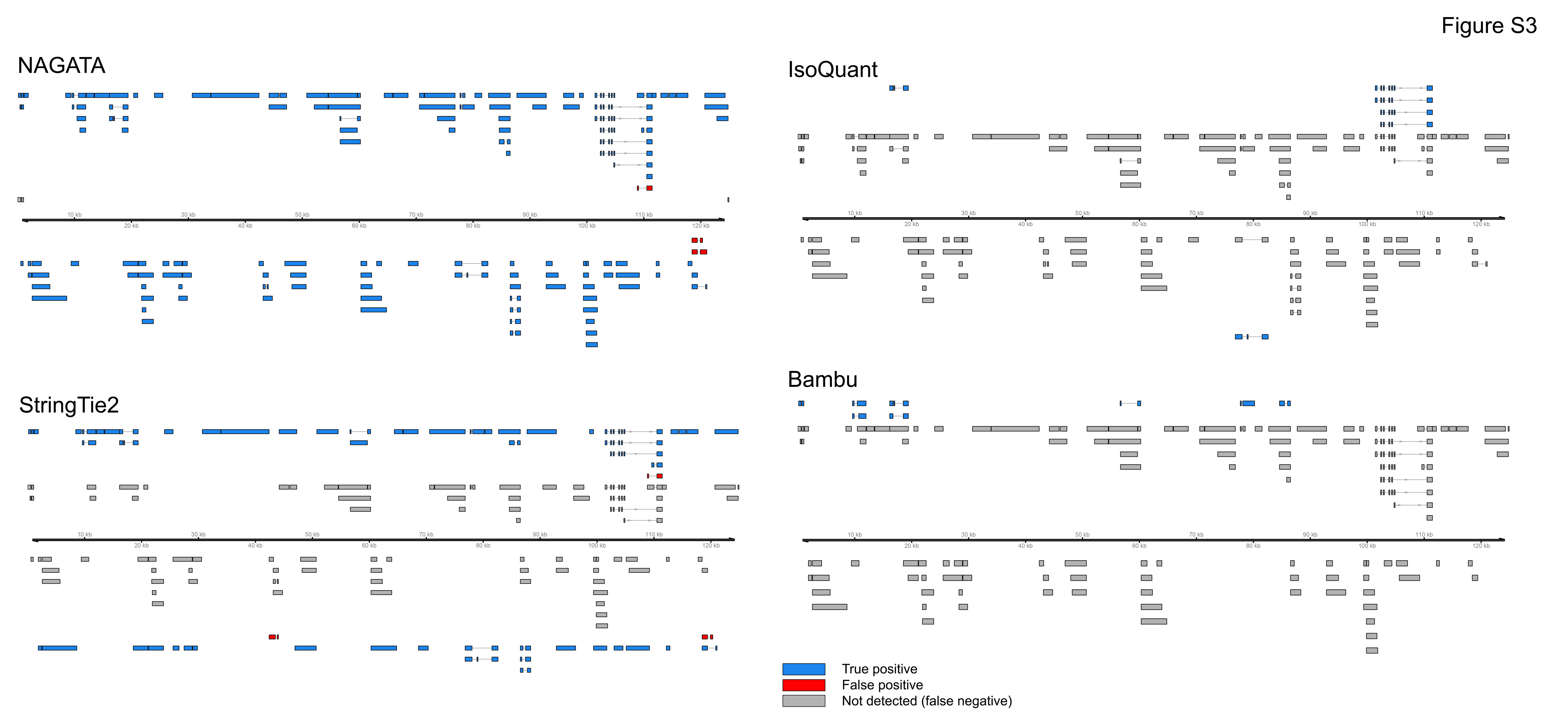

### Figure S4

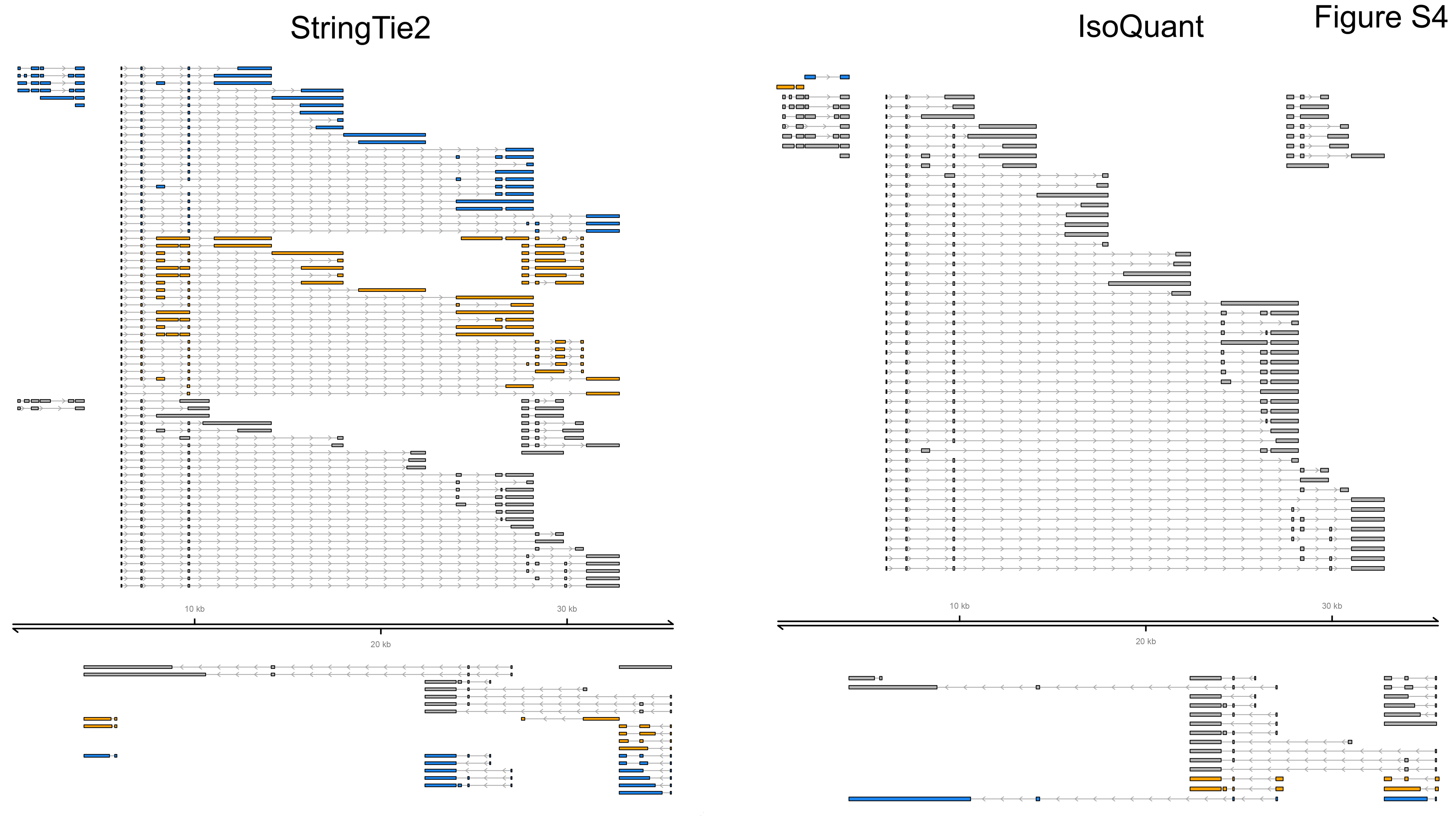

### Figure S5

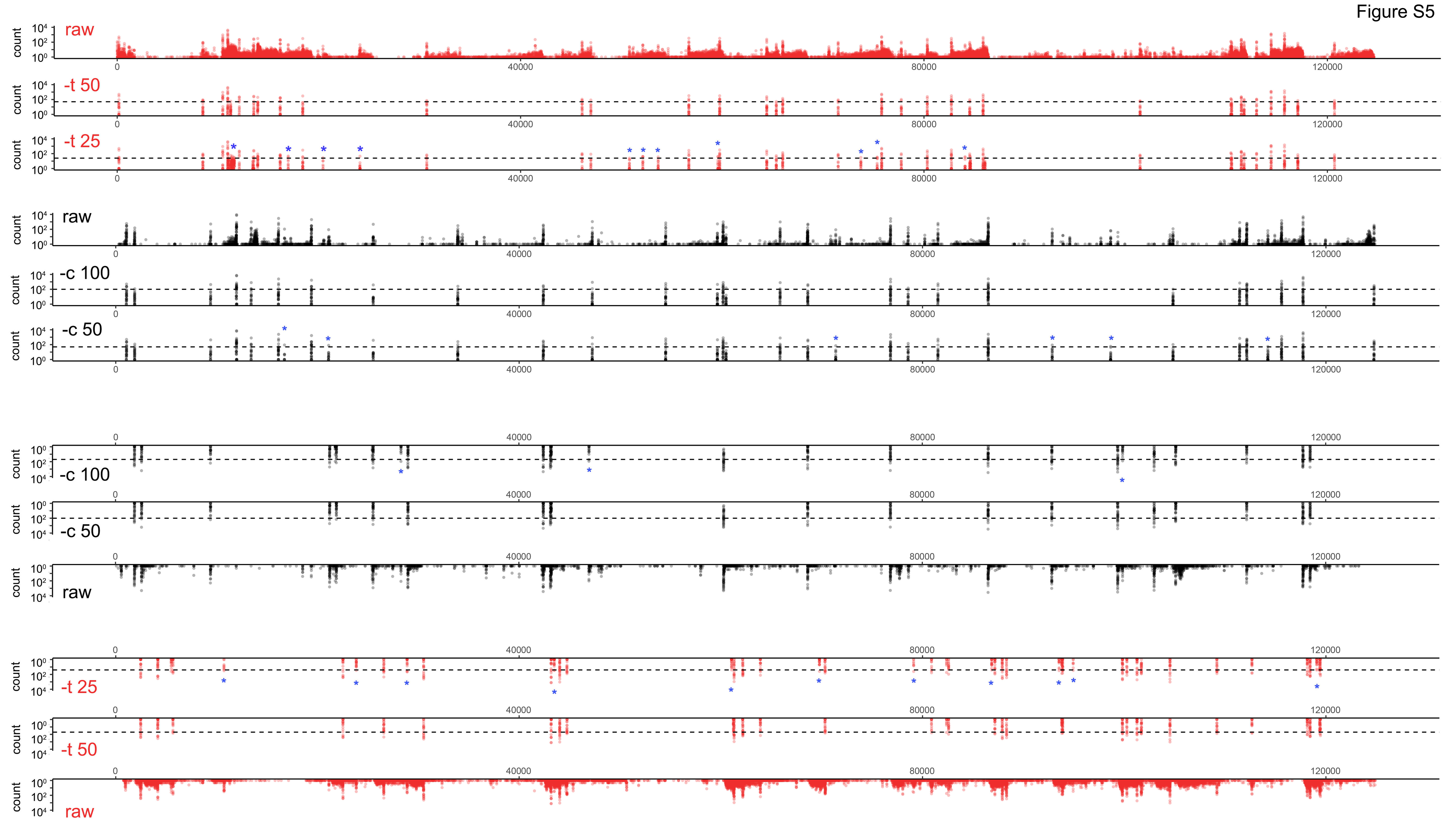

### Figure S6

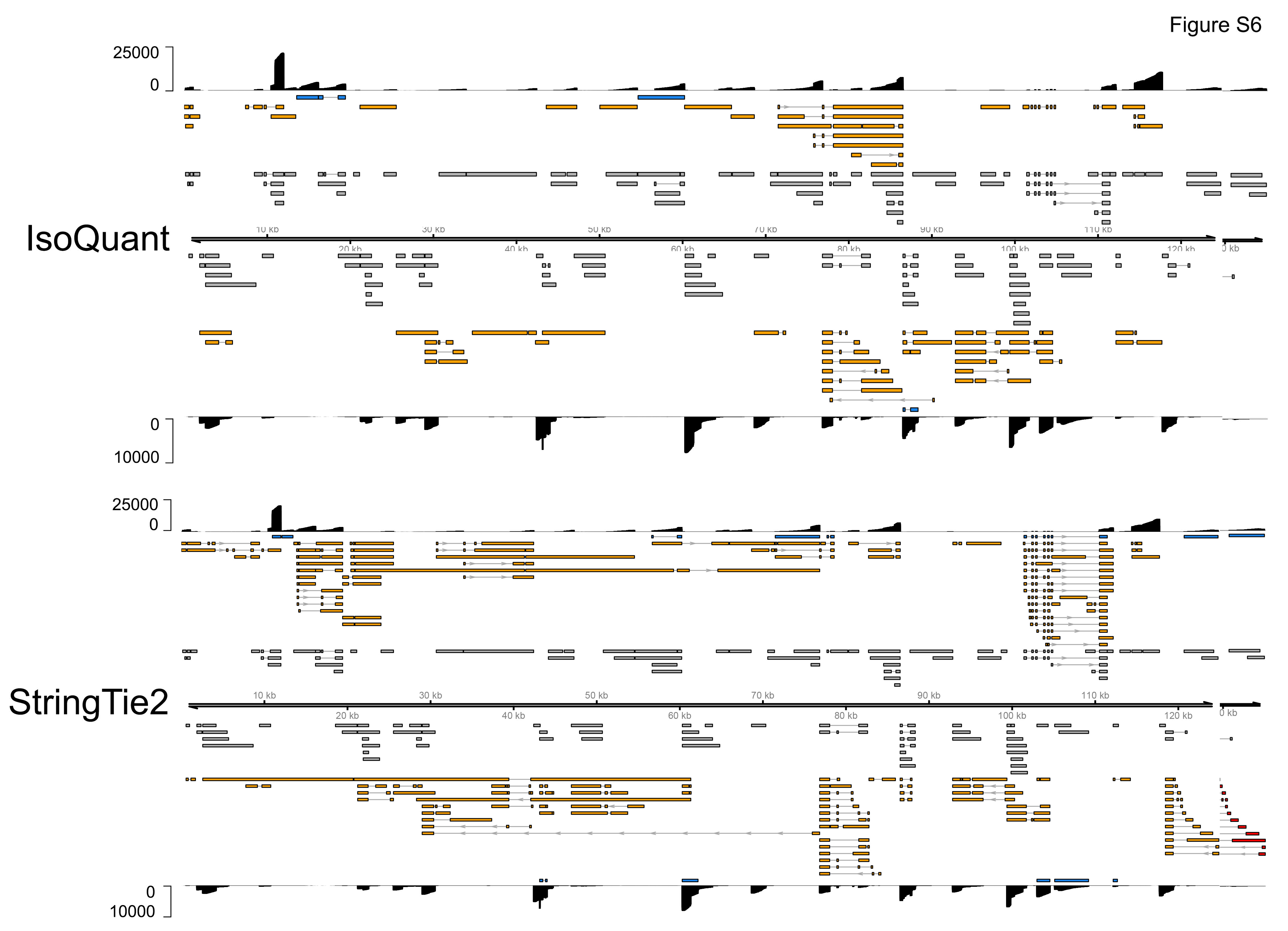
